# A rapid sporozoite viability assay identifies anti-*Cryptosporidium parvum* leads and targetable enzymatic activities in the invasive stage

**DOI:** 10.64898/2026.07.12.738121

**Authors:** Peng Jiang, Dongqiang Wang, Yiming Wang, Jigang Yin, Guan Zhu

## Abstract

Current phenotypic screens for anti-*Cryptosporidium* compounds typically quantify intracellular parasite growth in host cell cultures after two days of infection. Here, we developed a rapid, host cell-free phenotypic assay that directly measures compound-induced loss of viability in excysted *Cryptosporidium parvum* sporozoites, the invasive stage that initiates infection. We first compared qRT-PCR, luminescence ATP, and resazurin fluorescence readouts for detecting viable sporozoites. The luminescence ATP assay provided the best balance of linear dynamic range, assay time, parasite input, and cost, and was therefore adapted for high-throughput screening.

Screening 5,000 bioactive compounds at 40 μM identified 28 primary hits with ≥50% inhibition of sporozoite viability, including 14 with >60% inhibition. Secondary screening of these 14 compounds at 4 μM identified five hits retaining >50% inhibition: ZL0420, sulbactam, abexinostat, kojic acid, and SIB-1757. All five showed submicromolar activity against free sporozoites, with EC_50_ values of 0.073–0.311 μM. Four compounds, abexinostat, ZL0420, SIB-1757, and sulbactam, also inhibited intracellular parasite growth in vitro, with EC_50_ values of 0.316–11.87 μM and selectivity indices from >17 to >107. In an IFN-γ-knockout mouse model, these four compounds reduced oocyst shedding over the 35-day experiment by 53.7–79.0% based on area-under-the-curve analysis and improved body-weight trajectories and ileal histopathology.

Biochemical assays further showed that abexinostat inhibited native parasite HDAC activity at low nanomolar concentrations, while sulbactam inhibited a β-lactamase-like activity in sporozoite lysates. These findings establish sporozoite viability as a rapid screening endpoint and identify anti-*Cryptosporidium* leads associated with targetable enzymatic activities in the invasive stage.

**Author summary:** *Cryptosporidium parvum* is a major cause of diarrheal disease in humans and young animals, but treatment options remain limited. Most laboratory screens for new drugs against this parasite require infection of host cells and measurement of parasite growth after one or more days. We developed a faster approach that tests whether compounds can directly damage freshly excysted sporozoites — the parasite stage that first invades intestinal cells — or reduce their viability. This assay can be completed within a few hours and does not require host cells.

Using this approach, we screened 5,000 bioactive compounds and identified several molecules that rapidly reduced sporozoite viability. Four of these compounds also inhibited parasite growth in cell culture and reduced infection severity in a mouse model, as measured by parasite shedding, body-weight changes, and intestinal pathology. We further showed that one compound inhibits parasite histone deacetylase activity, while another inhibits a β-lactamase-like activity present in sporozoites. Our study provides a rapid screening strategy and highlights vulnerable biological activities in the invasive stage of *C. parvum*.

## Introduction

*Cryptosporidium* is a genus of globally distributed apicomplexan parasites, comprising more than 40 recognized species or genotypes that infect all major vertebrate groups [1]. Human cryptosporidiosis is caused primarily by the zoonotic species *Cryptosporidium parvum* and the anthroponotic species *C. hominis*, and may result in moderate to severe diarrheal disease [2,3]. Young children and immunocompromised individuals are among the populations most vulnerable to infection, in whom cryptosporidiosis is associated with substantial morbidity and mortality [4]. In farm animals, particularly in cattle, *C. parvum* is a major cause of neonatal diarrheal disease and can result in substantial economic losses [5–8]. Other *Cryptosporidium* species infect diverse mammals, birds, reptiles, amphibians, and fish, with implications for domestic animal health, wildlife conservation, and environmental transmission [1].

Despite its veterinary and medical importance, treatment options for cryptosporidiosis remain limited. Nitazoxanide is currently the only drug approved by the United States Food and Drug Administration to treat cryptosporidiosis in humans one year of age or older [4,9–11]. However, its efficacy is limited in immunocompromised patients and has not been established in infants, two populations at high risk for severe disease [12,13]. Other compounds, including paromomycin and clofazimine, have been evaluated experimentally or used off-label, but clinical and experimental outcomes have varied [14,15]. Approved options for treating active cryptosporidial infection in animals are also limited, although halofuginone lactate is licensed in the European Union and the United Kingdom primarily to reduce clinical signs and oocyst shedding in calves [16].

Over the past decade, increasing efforts have been devoted to anti-cryptosporidial drug discovery, leading to the identification of multiple preclinical leads with activity in cell culture and animal models [12,13]. Recent advances include development of controlled human infection models using current good manufacturing practice *C. parvum* oocysts under an investigational new drug application [17], as well as discovery and optimization of parasite-directed candidates such as phosphatidylinositol-4-OH kinase inhibitors [18]. Nonetheless, the absence of broadly effective and clinically approved anti-cryptosporidial drugs highlights the need for additional screening strategies, validated targets, and chemically diverse therapeutic leads.

Both phenotypic and target-based approaches have been used for anti-cryptosporidial drug discovery [13]. In most phenotypic screens, compounds are tested against intracellular parasite development in epithelial cell monolayers, and parasite burden is quantified after one to two days of infection. Three major assay formats are currently used for high-throughput or medium-throughput screening: luminescence assays using transgenic *C. parvum* expressing nanoluciferase, high-content imaging assays based on fluorescent labeling of the parasitophorous vacuole membrane, and qRT-PCR assays that quantify parasite RNA directly from infected cell lysates [19–22]. These approaches have been highly useful, but all require host cell culture and specific strategies to distinguish parasite-derived signals from host-cell background.

Here, we explored a complementary host cell-free approach that directly evaluates compound-induced loss of viability in excysted *C. parvum* sporozoites. Sporozoites are the motile invasive stage that emerges from oocysts and initiates infection of intestinal epithelial cells. Although free sporozoites survive only for a limited period outside host cells, we reasoned that this window may be sufficient to identify fast-acting compounds that compromise parasite viability before invasion. Such compounds would be expected to enrich for inhibitors of biological processes that are essential in the invasive stage and may also affect subsequent parasite development in vitro and in vivo.

We first compared qRT-PCR, luminescence ATP, and resazurin fluorescence assays for quantifying viable sporozoites and selected the luminescence ATP assay for screening because it provided a favorable balance of dynamic range, sensitivity, speed, and cost. Using this assay, we screened 5,000 bioactive compounds and identified five hits with submicromolar activity against free sporozoites. Four of these compounds also inhibited intracellular parasite growth in vitro and showed efficacy in an IFN-γ-knockout mouse infection model. Finally, we performed biochemical assays supporting inhibition of native parasite HDAC activity by abexinostat and inhibition of a β-lactamase-like activity in sporozoite lysates by sulbactam. Together, these findings establish sporozoite viability as a rapid phenotypic screening endpoint and identify chemically diverse anti-*Cryptosporidium* leads associated with targetable enzymatic activities in the invasive stage.

## Materials and methods

### Ethics statement

Bovine calves were used for propagation of Cryptosporidium parvum oocysts. Interferon-γ-knockout mice were used to evaluate the anti-cryptosporidial efficacy of selected compounds identified in this study. All animal procedures complied with the Guide for the Care and Use of Laboratory Animals of the Ministry of Health of China. Animal use protocols were approved by the Animal Welfare and Research Ethics Committee of the Institute of Zoonosis, Jilin University (AUP #2020-IZ-20 and #2024-IZ-00031). Mice were monitored for body weight and general health, including physical activity, fur condition, body posture, and mental state.

### Parasite propagation and preparation of excysted sporozoites

*C. parvum* oocysts, genotype IIaA17G2R1 at the gp60 locus, were propagated in calves. Oocysts were purified by Sheather’s sucrose flotation and stored at 4 °C in PBS containing 200 units/mL penicillin and 0.2 mg/mL streptomycin, as described previously [23–25]. Before quantitative experiments, oocysts were suspended in water and treated with 0.2% NaOCl on ice for 5 min, followed by five to eight washes in PBS.

For excystation, oocysts were incubated in RPMI-1640 medium containing 0.75% taurodeoxycholic acid at 37 °C for 45–60 min, followed by three washes with PBS [26]. Purified sporozoites were counted with a hemocytometer and used immediately for subsequent experiments. Oocysts used in this study were less than 3 months old and had excystation rates >80%.

### Evaluation of three sporozoite viability assays

Three assays were compared for their ability to quantify viable *C. parvum* sporozoites in a host cell-free format: qRT-PCR, luminescence ATP assay, and resazurin-based fluorescence assay. In all assays, freshly excysted sporozoites were suspended in RNase-/protease-free PBS (pH 7.4; Beyotime Biotechnology, Shanghai, China) containing 1% bovine serum albumin (BSA; Yeasen Biotechnology, Shanghai, China). Unless otherwise specified, sporozoites were incubated at 25 °C for 3 h before assay readout.

To determine the extracellular assay window, sporozoites were incubated at 25 °C for 1–12 h and then analyzed by qRT-PCR and luminescence ATP assays. For qRT-PCR, sporozoite suspensions were centrifuged at 12,000 × g for 5 min, and pellets were processed for RNA isolation. For the luminescence ATP assay, sporozoites were processed directly for ATP measurement. After determining that sporozoites retained near-baseline viability for up to approximately 4 h under these conditions, a 3 h incubation period was used for subsequent assay comparison, high-throughput screening, and dose-response assays.

The qRT-PCR assay quantified *C. parvum* 18S rRNA, as described previously [21,27]. Total RNA was isolated from sporozoites using the RNeasy Mini Kit (Qiagen, Venlo, Netherlands; Cat. #74104). The primers were Cp18S-1011F (5′-TTG TTC CTT ACT CCT TCA GCA C-3′) and Cp18S-1185R (5′-TCC TTC CTA TGT CTG GAC CTG-3′). Reactions were performed using HiScript II One-Step qRT-PCR SYBR Green Kit (Vazyme, Nanjing, China) on an Applied Biosystems QuantStudio 1 real-time PCR system (Thermo Fisher Scientific, Carlsbad, CA, USA). Cycling threshold (C_T_) values were recorded and plotted against the number of sporozoites on a logarithmic scale.

The luminescence ATP assay was performed using the BacTiter-Lumi Luminescent Microbial Cell Viability Assay Kit (Beyotime Biotechnology), which detects ATP through a luciferase-catalyzed reaction between ATP and luciferin. Briefly, 100 μL of sporozoite suspension was added to black-walled, flat-bottom 96-well plates (Labselect brand, Labgic Technology, Beijing, China), followed by addition of 100 μL BacTiter-Lumi reagent. Plates were shaken for 2 min and incubated for 10 min before luminescence signals were measured using a Cytation 5 Cell Imaging Multi-Mode Reader (BioTek Instruments, Winooski, VT, USA). Because this assay was selected for screening DMSO-dissolved compounds, the effects of DMSO concentration, freeze-thaw cycles, shaking time, and post-reaction incubation time were further evaluated.

The fluorescence assay measured NAD(P)H-dependent reduction of resazurin to resorufin [28]. It has been adapted to evaluating viability in *Plasmodium*, *Trypanosoma*, and *Leishmania* parasites [29–31]. Resazurin (MedChemExpress, New Jersey, USA) was mixed with sporozoites at a final concentration of 30 μg/mL in 100 μL of 1% BSA/PBS in 96-well microplates. The mixtures were incubated at 25 °C for 3 h, and fluorescence signals were measured using a Cytation 5 Cell Imaging Multi-Mode Reader (BioTek Instruments) with excitation at 560 nm and emission at 590 nm.

### High-throughput screening of compounds against sporozoite viability

A library of 5,000 bioactive compounds (Topscience, Shanghai, China) was screened for activity against *C. parvum* sporozoite viability using the luminescence ATP assay. Compounds were supplied as DMSO stocks and screened in black-walled 96-well microplates. Each well contained 5 × 10^5^ freshly excysted sporozoites in 100 μL PBS/1% BSA and compound at a final concentration of 40 μM. After incubation at 25 °C for 3 h, 100 μL BacTiter-Lumi reagent was added to each well. Plates were shaken for 2 min, incubated for 8–10 min, and read using the Cytation 5 Cell Imaging Multi-Mode Reader. All compounds were evaluated in duplicate, and all wells contained a final DMSO concentration of 0.5%.

Each screening plate included blank controls containing reaction buffer only, DMSO negative controls representing maximal sporozoite viability, and H_2_O_2_ positive controls representing minimal viability. In the current assay setup, the positive-control wells contained 1 mM H_2_O_2_ and 0.5% DMSO. Percent inhibition of sporozoite viability was calculated as:

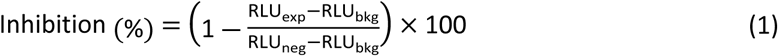

where *RLU_exp_*, *RLU_neg_* and *RLU_bkg_* denote relative luminescence units from compound-treated wells, DMSO-treated negative-control wells, and blank wells, respectively.

Assay performance was evaluated by calculating a conservative SD-based signal window (SW) and the Z′ factor, as described for HTS assay validation [32–35]. The SW was calculated as:

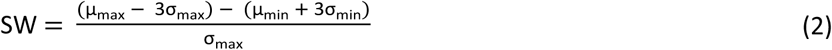

And the Z’ factor was calculated as:

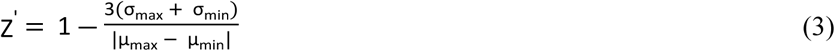

where *μ_max_* and *σ_max_* are the mean and standard deviation of the maximal signal controls, i.e., DMSO-treated sporozoites, and *μ_min_* and *σ_min_* are the mean and standard deviation of the minimal signal controls, i.e., H_2_O_2_-treated sporozoites.

Compounds showing >60% inhibition in the primary screen were evaluated in a secondary screen at 4 μM using the same luminescence ATP assay. Hits retaining >50% inhibition at 4 μM were selected for dose-response analysis.

### Dose-response assays and morphological assessment of sporozoites

Selected hits were evaluated in dose-response assays to determine 50% effective concentrations (EC_50_ values) against free sporozoites. Freshly excysted *C. parvum* sporozoites (5 × 10^5^/well) were treated with serially diluted compounds in 1% BSA/PBS containing 0.5% DMSO. After incubation at 25 °C for 3 h, sporozoite viability was measured using the luminescence ATP assay as described above. Percent inhibition values were plotted against compound concentrations, and EC_50_ values were calculated by nonlinear regression using a four-parameter logistic model, with the lower plateau constrained to zero unless otherwise specified:

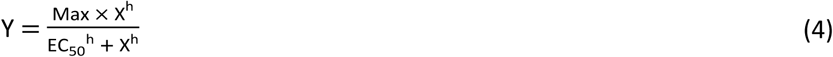

where *Y* is the percent inhibition, *Max* is the maximal inhibition, *X* is inhibitor concentration, *EC*_50_ is concentration producing 50% inhibition, and *h* is the Hill’s slope.

The effects of selected hits on sporozoite morphology were evaluated by immunofluorescence microscopy. Excysted sporozoites were treated with compounds at 10 μM or with diluent control (0.5% DMSO) in 1% BSA/PBS for 10, 30, 60, or 120 min. Samples were fixed with 4% paraformaldehyde, permeabilized with 0.5% Triton X-100, blocked in 5% nonfat dry milk, and labeled with a homemade rabbit polyclonal antibody against a C. parvum membrane protein.

Samples were then labeled with Alexa Fluor 488-conjugated goat anti-rabbit IgG secondary antibody, counterstained with 4′,6-diamidino-2-phenylindole (DAPI; Sigma-Aldrich, St. Louis, MO, USA; 1.0 μg/mL), mounted with antifade mounting medium (Beyotime Biotechnology), and examined using an Olympus BX53 research microscope (Olympus/Evident, Tokyo, Japan). All washes and dilutions were performed in PBS, and procedures were performed at room temperature unless otherwise specified.

### In vitro anti-cryptosporidial efficacy and host-cell cytotoxicity assays

Selected hits were evaluated for inhibition of intracellular *C. parvum* growth using an established 44 h infection assay with parasite burden quantified by qRT-PCR [21,22,27]. HCT-8 cells, a human ileocecal adenocarcinoma cell line (ATCC #CCL-244), were seeded into 96-well microplates and cultured in RPMI-1640 medium supplemented with 10% fetal bovine serum (FBS) at 37 °C until reaching approximately 80% confluence.

Cell monolayers were inoculated with *C. parvum* oocysts (2 × 10^4^ oocysts/well) in culture medium containing 0.15% taurocholic acid. After incubation at 37 °C for 3 h to allow excystation and invasion, uninvaded parasites and debris were removed by medium exchange. Test compounds at specified concentrations were then added in fresh culture medium. Each plate included blank controls, diluent controls containing 0.5% DMSO, and positive controls containing 2 μM nitazoxanide. All wells contained a final DMSO concentration of 0.5%.

Infected cells were cultured for an additional 41 h at 37 °C, corresponding to a total infection time of 44 h. At this time point, most parasites had developed into mature meronts, with some parasites entering sexual stages. Plates were centrifuged at 1,000 × g for 10 min to secure cell monolayers. After supernatants were removed, 100 μL ice-cold 0.1% BSA prepared in nuclease-/proteinase-free water was added to each well. Plates were placed in an ice bucket and shaken in a multi-tube vortexer (YKX-200, Yika Technology, Shanghai, China) for 30 min. Plates were then centrifuged at 2,000 × g for 10 min at 4 °C, and supernatants were collected for qRT-PCR analysis.

qRT-PCR was performed using HiScript II One-Step qRT-PCR SYBR Green Kit (Vazyme, Nanjing, China). Reactions were performed in 96-well PCR plates with 20 μL reaction volume, containing 2 μL diluted cell lysate (1:100 dilution) and primers Cp18S-1011F and Cp18S-1185R for *C. parvum* 18S rRNA. Host-cell 18S rRNA was quantified for normalization using primers Hs18S-1F (5′-GGC GCC CCC TCG ATG CTC TTA-3′) and Hs18S-1R (5′-CCC CCG GCC GTC CCT CTT A-3′). Thermal cycling was performed as described above for the qRT-PCR-based sporozoite assay. Relative parasite burden was calculated using the 2^-ΔΔC_T_ method [22], normalized to host-cell 18S rRNA, and percent inhibition of intracellular parasite growth was calculated relative to DMSO-treated controls. EC_50_ values were calculated by nonlinear regression using the four-parameter logistic model described above.

Host-cell cytotoxicity was evaluated in HCT-8 cells using a Cell Proliferation Assay Kit (MTS; Saint-Bio Biotechnology, Shanghai, China) according to the manufacturer’s instructions. Briefly, HCT-8 cells were seeded in 96-well plates at 5 × 10^3^ cells/well and cultured for 24 h. Cells were then treated with selected compounds at indicated concentrations or with diluent control (0.5% DMSO) for 44 h at 37 °C. Culture medium was discarded, and monolayers were gently rinsed once with PBS. MTS reagent was added at 20 μL/well, followed by incubation for 1 h at 37 °C. Absorbance at 490 nm was measured using a Synergy LX multimode plate reader (BioTek Instruments, Winooski, VT, USA). The 50% toxic concentrations (TC_50_ values) were determined by nonlinear regression using GraphPad Prism version 10 or higher (GraphPad Software, Boston, MA, USA). In vitro selectivity indices (SI) were calculated as TC_50_ against HCT-8 cells divided by EC_50_ against intracellular parasite growth (SI = TC_50_/EC_50_).

### In vivo efficacy in IFN-γ-knockout mice

Four compounds with confirmed in vitro activity were evaluated for in vivo efficacy using an IFN-γ-KO mouse model of *C. parvum* infection [36–38]. Female IFN-γ-KO mice, 6–8 weeks old, were infected by oral gavage with 5 × 10^4^ transgenic *C. parvum* oocysts expressing nanoluciferase in 200 μL PBS (n = 6 mice/group). The transgenic parasite was generated by insertion of an mNeonGreen-Nluc-Neo cassette into the thymidine kinase locus, as described previously [19].

Beginning at 3 days post-infection (dpi), mice received once-daily treatment for seven consecutive days. Abexinostat was administered by oral gavage at 25 mg/kg/day. ZL0420, sulbactam sodium, and SIB-1757 were administered by intraperitoneal injection at 10, 20, and 20 mg/kg/day, respectively. Compounds were formulated in vehicle containing 10% DMSO, 40% PEG300, 5% Tween-80, and 45% saline. Vehicle-control mice received vehicle only once daily. Mice were monitored at least once daily for body weight, activity, fur condition, posture, and mental state; animals showing severe distress or reaching protocol-defined humane endpoints were to be humanely euthanized.

Before efficacy testing, selected doses were evaluated for short-term tolerability in uninfected female C57BL/6 mice, 6–8 weeks old, three mice per dose group, using once-daily dosing for seven days. Body weight and general health were monitored daily. Health scoring included physical activity, fur condition, body posture, and mental state.

During the 7-day treatment period in infected IFN-γ-KO mice, body weights and fecal samples were collected daily. After treatment, body-weight and fecal-sample collection continued every other day until 21 dpi and then once weekly until 35 dpi, the final day of the experiment. On 35 dpi, mice were deeply anesthetized by intraperitoneal injection of pentobarbital sodium (50 mg/kg), followed after 5–10 min by cervical dislocation to ensure death, and intestinal tissues were collected for histopathology.

Nanoluciferase activity in fecal samples was measured using the Nano-Glo Luciferase Assay Kit (Promega, Madison, WI, USA), following the manufacturer’s instructions unless otherwise specified. Briefly, 20 mg fecal material was placed into a 1.5 mL microtube containing 0.5 mL lysis buffer (50 mM Tris, pH 8.0; 2 mM EDTA; 2 mM DTT; 1% Triton X-100; 10% glycerol in sterile water) and 3 mm ceramic beads (Beyotime Biotechnology). Samples were homogenized by vortexing for 5 min using a tissue homogenizer and centrifuged at 8,000 × g for 1 min [39]. Three 100 μL aliquots of supernatant, each corresponding to 4 mg feces, were transferred into white 96-well plates. Nano-Glo substrate reconstituted with Nano-Glo buffer at a 1:50 dilution was added at 100 μL/well. Luminescence was measured immediately using the Cytation 5 Cell Imaging Multi-Mode Reader, and data were expressed as relative luminescence units (RLU) per 4 mg feces.

### Histopathology and morphometric analysis

Ileal tissues collected at 35 dpi were processed into 1–2 cm segments, rinsed with cold PBS to remove intestinal contents and debris, fixed in 10% neutral-buffered formalin, paraffin-embedded, and sectioned at 5 μm thickness. Tissue sections were deparaffinized with xylene, rehydrated through a graded ethanol series (100%, 95%, 85%, and 75%), rinsed with distilled water, stained with hematoxylin and eosin (H&E), dehydrated, cleared, and mounted with neutral resin according to standard histopathology protocols. Slides were examined and digitized using a SLIDEVIEW VS200 slide scanner (Evident/Olympus, Tokyo, Japan).

For morphometric analysis, ileal tissues from three mice per group were examined. For each mouse, ten well-oriented villus–crypt units were selected from H&E-stained digital images, yielding 30 villus–crypt measurements per group. Villus length was measured from the villus tip to the villus–crypt junction, and crypt depth was measured from the villus–crypt junction to the base of the crypt. The villus length/crypt depth ratio was calculated for each measured unit. Because multiple villus–crypt units were measured from each mouse, statistical comparisons were performed using nested one-way ANOVA, with mouse nested within treatment group, followed by Dunnett-adjusted multiple comparisons.

### Bioinformatic and proteomic analyses of candidate enzymatic activities

Annotated *C. parvum* proteins potentially related to the known activities of selected compounds were identified by keyword search and domain-based annotation in CryptoDB, together with BLASTP searches when appropriate. Candidate HDAC/Sir2-family proteins, metallo-β-lactamase or metallo-β-lactamase domain-containing proteins, bromodomain-containing proteins, and mGluR5-like proteins were examined.

For CpHDAC abundance in sporozoites, spectral-count information was obtained from available CryptoDB proteomic evidence derived from a previously published *C. parvum* sporozoite proteome dataset [40]. Quantitative protein abundance values were obtained from our recently published DIA-MS-based *C. parvum* sporozoite proteome/phosphoproteome dataset [41]. Protein abundance from the latter dataset was expressed as log_2_-transformed abundance values. No new mass spectrometry data were generated for this analysis.

### Biochemical assays for HDAC and β-lactamase-like activities

Two selected compounds were evaluated for inhibition of candidate native enzymatic activities in *C. parvum* sporozoite lysates: abexinostat against HDAC activity and sulbactam against β-lactamase-like activity. Sporozoite lysates were prepared using NP-40 lysis buffer (Thermo Fisher Scientific Inc., Waltham, MA, USA) containing protease inhibitor cocktail (MedChemExpress, New Jersey, USA). To prepare parasite lysates, 1 × 10^9^ freshly excysted sporozoites were suspended in 75 μL lysis buffer and subjected to five freeze-thaw cycles between liquid nitrogen and ice. Samples were then incubated on ice for 20 min and centrifuged at 16,000 × g. Supernatants were collected, and protein concentrations were quantified by bicinchoninic acid (BCA) assay using BSA as the standard.

For HDAC assays, HCT-8 cell lysates were prepared similarly for comparison, using 2 × 10^6^ cells in 100 μL lysis buffer. For β-lactamase assays, *Escherichia coli* BL21 lysates were prepared using the same NP-40-based extraction procedure to maintain comparable lysis conditions across samples and preserve native enzymatic activity.

HDAC activity was measured using an Amplite Fluorimetric HDAC Activity Assay Kit, green fluorescence (AAT Bioquest, Pleasanton, CA, USA). Reactions were performed in 100 μL volume in 96-well fluorescence microplates. Each reaction contained 40 μL parasite or host-cell lysate diluted in assay buffer to provide 5 μg total protein, 50 μL HDAC Green substrate working solution, and abexinostat or vorinostat at specified concentrations. The HDAC Green substrate working solution was freshly prepared in assay buffer by mixing HDAC Green substrate, signal enhancer, and assay buffer at a ratio of 1:5:250. Test compounds were diluted in assay buffer, and the final DMSO concentration was maintained at 0.5% in all reactions. After incubation at 25 °C for 45 min, fluorescence was measured using a Cytation 5 Cell Imaging Multi-Mode Reader with excitation at 490 nm and emission at 525 nm.

β-Lactamase-like activity was assayed using nitrocefin as the substrate [42]. In addition to parasite and bacterial lysates, recombinant *E. coli* β-lactamase purchased from Aladdin Biochemical Technology (Shanghai, China) was included as a positive control. Reactions were performed in a final volume of 100 μL/well, containing 50 μL PBS supplemented with 30 μU recombinant β-lactamase or 30 μg parasite or bacterial total protein, together with 0.5% DMSO or sulbactam at specified concentrations. After incubation at 25 °C for 30 min, 50 μL nitrocefin solution (0.1 mg/mL in PBS) was added. Reactions were incubated for an additional 20 min at 25 °C, and optical density was measured at 490 and 380 nm. Relative β-lactamase-like activity was determined from the OD490/OD380 ratio.

HDAC and β-lactamase assays were performed in triplicate reactions. Blank controls containing buffer only were included for background subtraction. Percent inhibition was calculated from background-subtracted signals in reactions with and without inhibitors, using the same general formula described above for the luminescence ATP assay. Dose-response curves were fitted using the four-parameter logistic model described above or, when curve asymmetry was evident, a five-parameter logistic model [43].

### Data analysis and statistics

Data were analyzed using GraphPad Prism version 10 or higher (GraphPad Software, Boston, MA, USA). Unless otherwise specified, data are presented as mean ± standard error of the mean (SEM). HTS assays were performed in duplicate wells, while dose-response, cytotoxicity, and biochemical assays were performed in triplicate unless otherwise indicated. All compound-treated reactions and their corresponding controls contained 0.5% DMSO.

Linear regression was used to determine the linear dynamic range of sporozoite detection assays. Pearson correlation analysis was used to compare qRT-PCR and luminescence ATP readouts. EC_50_, TC_50_, and IC_50_ values were determined by nonlinear regression using four-parameter logistic or five-parameter logistic models, as appropriate.

For mouse infection experiments, fecal oocyst shedding was expressed as RLU per 4 mg feces. Area under the curve (AUC) values for oocyst shedding were calculated using the trapezoidal method for the whole observation period (3–35 dpi), treatment period (3–9 dpi), and post-treatment period (9–35 dpi). Relative oocyst shedding was calculated as AUC of the treatment group divided by AUC of the vehicle group, and percent reduction was calculated as 1 – AUC_(treatment)_/AUC_(vehicle)_. Survival curves were compared using the log-rank test. Oocyst shedding and body-weight changes were compared among groups at individual time points using one-way ANOVA followed by Dunnett’s multiple comparison test against the indicated control group. Morphometric measurements were analyzed using nested one-way ANOVA, with mouse nested within treatment group, followed by Dunnett-adjusted multiple comparisons. P values <0.05 were considered statistically significant.

## Results

### Comparison of three *C. parvum* sporozoite viability assays for performance and cost-effectiveness

Unlike assays that quantify intracellularly developing *C. parvum* in host cell monolayers, assays using free sporozoites can directly monitor parasite viability without the need to distinguish parasite-derived signals from host-cell background. Among available viability readouts, we compared three assays for performance, operational time, parasite input, and cost-effectiveness: an established qRT-PCR assay that quantifies parasite 18S rRNA [21,27], a luminescence ATP assay [44,45], and an NAD(P)H-dependent resazurin fluorescence assay [28].

Freshly excysted sporozoites were suspended in 1% BSA/PBS and incubated at 25 °C for 3 h unless otherwise specified. The lower incubation temperature was used to slow sporozoite metabolism and extend the host cell-free assay window, while BSA was included to reduce potential colloidal aggregation effects of small molecules [46]. Under these conditions, sporozoites retained near-baseline viability for up to 4 h, as determined by both qRT-PCR and luminescence ATP assays, and showed no obvious morphological deterioration during the first 3 h (Fig. 1A–C). Based on these observations, a 3 h incubation period was used for subsequent assay comparison, screening, and dose-response experiments.

**Fig 1.**
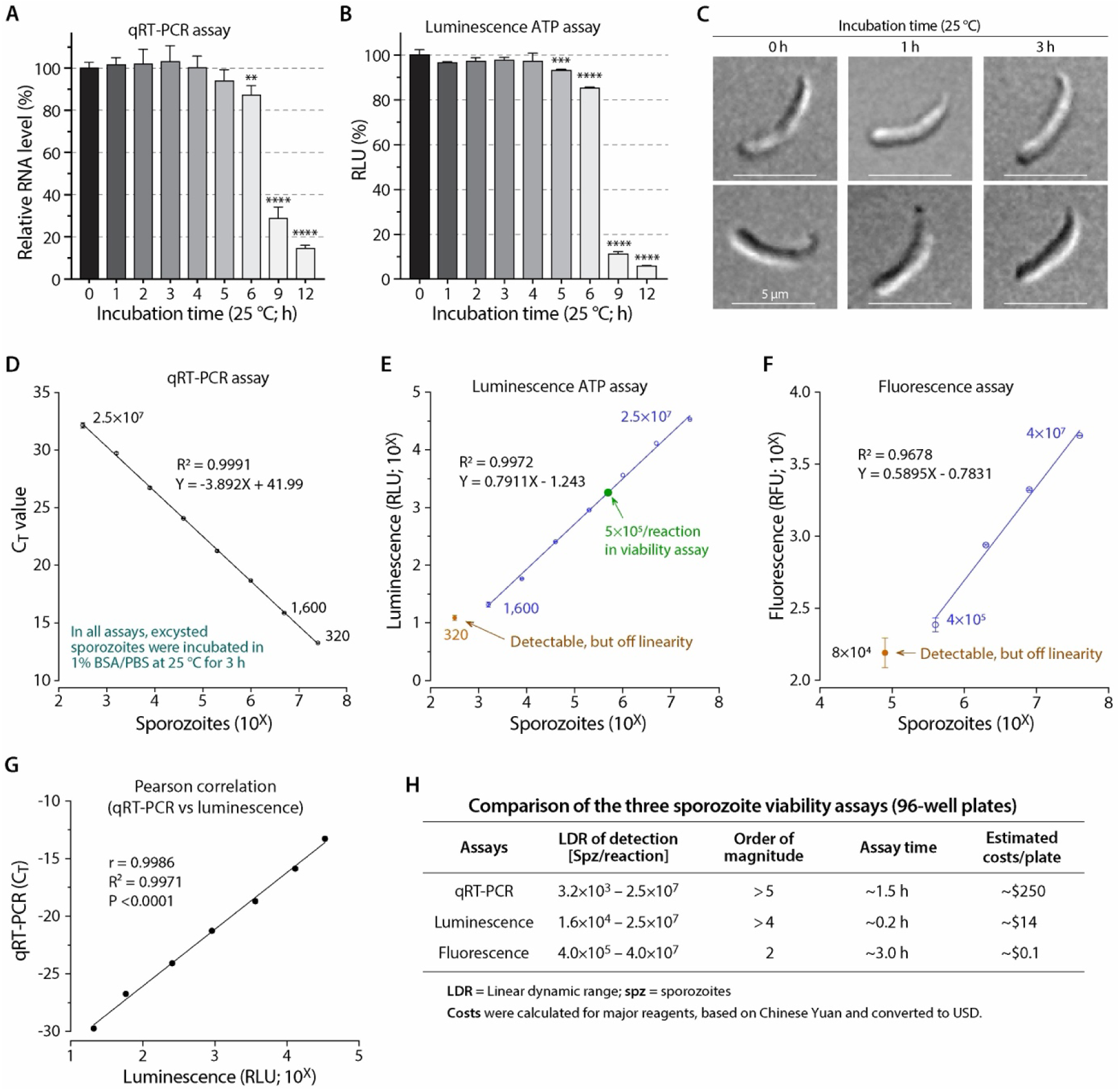
Performance comparison of three assays in the evaluation of the viability of post-excystation sporozoites of *C. parvum*. (A–B) Viability of post-excystation sporozoites incubated at 25 °C for various time periods as determined by qRT-PCR (**A**) and luminescence assay (**B**) assays, which quantified parasite 18S rRNA and ATP levels, respectively. RLU, relative luminescence unit. (**C**) Differential interference microscopy showing normal morphology of post-excystation sporozoites after incubation at 25 °C for up to 3 h. (**D–F**) Linear dynamic ranges of qRT-PCR assay (**D**), luminescence ATP assay (**E**), and resazurin-based fluorescence assay (**F**) in the detection of post-excystation sporozoites. (**G**) qRT-PCR and luminescence assays displaying excellent Pearson correlation (r = 0.9986, R^2^ = 0.9971, P <0.0001). (**H**) Comparison of performance parameters between qRT-PCR, luminescence, and fluorescence assays, including linear dynamic range (LDR), approximate assay time, and estimated costs. In all assays, sporozoites were prepared by excystation in RPMI-1640 medium containing 0.75% taurodeoxycholic acid. Post-excystation sporozoites were suspended in PBS/1% BSA, incubated at 25 °C for specified time periods before assays.

Among the three assays, qRT-PCR showed the broadest linear dynamic range, with >5 orders of magnitude of linearity, followed by the luminescence ATP assay with >4 orders of magnitude. The resazurin fluorescence assay showed the narrowest linear dynamic range, covering approximately 2 orders of magnitude and requiring substantially more sporozoites per reaction (Fig. 1D–F). Although qRT-PCR was the most analytically sensitive assay, luminescence ATP signals showed excellent correlation with qRT-PCR readouts within the linear range (Pearson’s r = 0.9986, R^2^ = 0.9971, P < 0.0001) (Fig. 1G).

Operational considerations favored the luminescence ATP assay for high-throughput screening. qRT-PCR provided the best linearity and sensitivity, but was more expensive and required RNA extraction and amplification. The resazurin fluorescence assay was the least expensive, but required a longer incubation time and a higher sporozoite input. The luminescence ATP assay provided a practical balance of dynamic range, assay time, parasite requirement, and cost, and was therefore selected for screening compounds against sporozoite viability (Fig. 1H).

We further optimized the luminescence ATP assay for high-throughput format. Freeze-thaw cycles increased luminescence signals from both oocyst and sporozoite samples, but non-freeze-thawed sporozoites produced sufficiently strong signals; this step was therefore omitted to streamline the assay (S1A Fig). Shaking for 1–4 min had little effect on signal intensity (S1B Fig), and DMSO concentrations up to 2% did not substantially affect luminescence signals (S1C Fig). After addition of the luminescence reagent, signals increased slightly during the first 10 min and then gradually declined, remaining suitable for measurement during the early post-reaction window (S1D Fig). In samples containing defined mixtures of live and ATP-depleted sporozoites, the assay showed excellent linearity between luminescence signal and the proportion of live sporozoites (R^2^ = 0.9957; S1E Fig).

Based on these results, the standard luminescence ATP assay consisted of freshly excysted sporozoites suspended in 1% BSA/PBS, incubation with compounds or diluent at 25 °C for 3 h in 96-well luminescence plates, addition of BacTiter-Lumi reagent, shaking for 2 min, incubation for 8 min, and luminescence measurement. All reactions contained 0.5% DMSO and were performed in duplicate for screening or triplicate for dose-response assays.

### High-throughput screening identifies five compounds with submicromolar activity against sporozoites

We next adapted the luminescence ATP assay for high-throughput screening of a 5,000-compound bioactive library. The screening and validation workflow included primary screening at 40 μM, secondary screening of selected hits at 4 μM, dose-response assays against free sporozoites, evaluation of intracellular parasite growth and host-cell cytotoxicity, and in vivo efficacy testing of validated hits (Fig. 2A).

**Fig 2.**
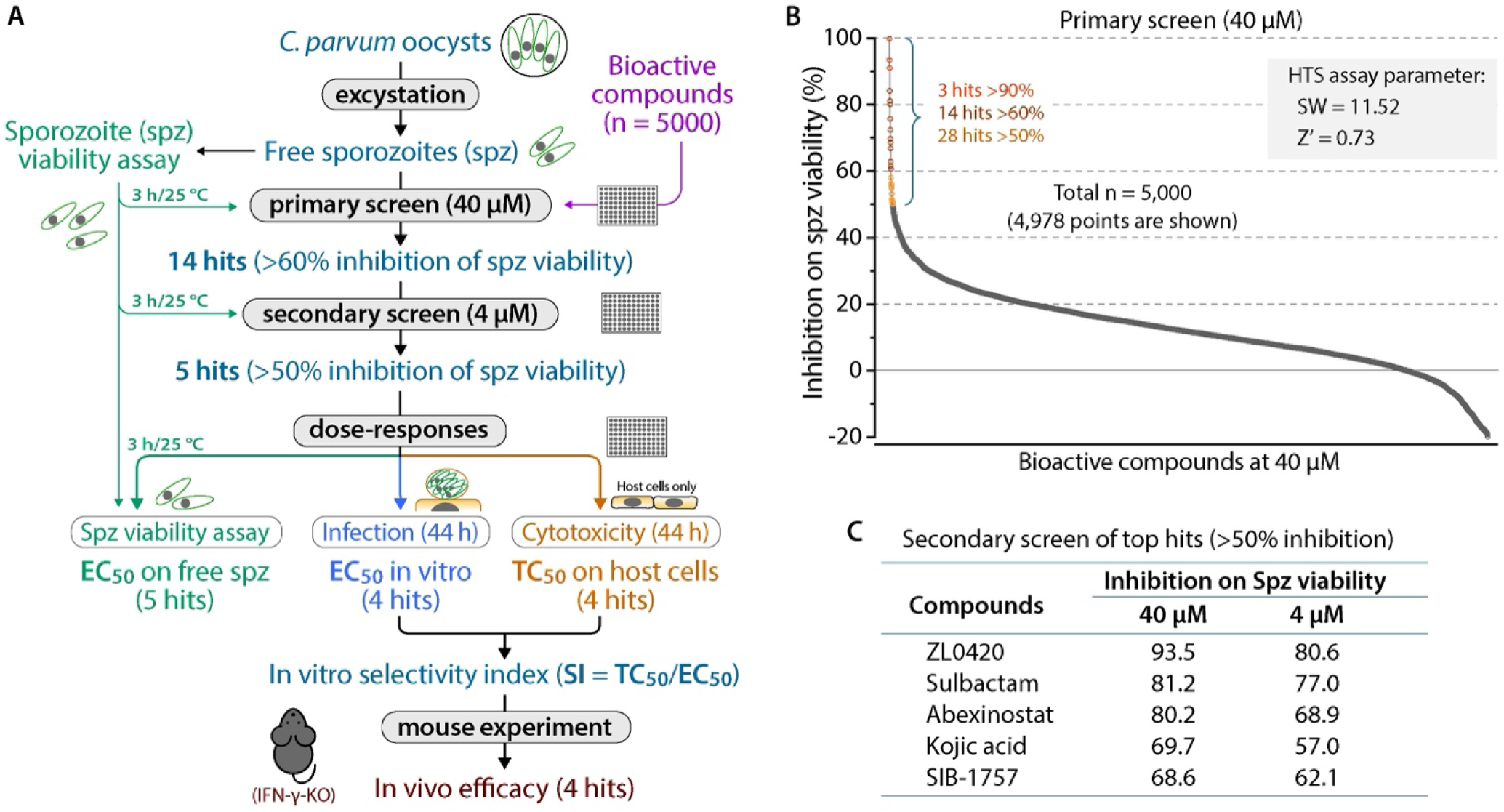
High-throughput screening of 5,000 bioactive compounds against the viability of *C. parvum* post-excystation sporozoites with luminescence ATP assay. (**A**) Schematic of the overall experimental design, including excystation, primary and secondary screening, evaluation of hits for in vitro efficacies on sporozoite viability and parasite intracellular development, and in vivo efficacies in a mouse infection model. (**B**) Summary of primary high-throughput screening (HTS) of 5,000 bioactive compounds at 40 μM using luminescence-based sporozoite viability assay. Signal window (SW) and Z’-factor of the assay are indicated. (**C**) Summary of five top hits identified by secondary screening at 4 μM. Spz, sporozoites.

Assay performance was evaluated using DMSO-treated sporozoites as the maximal signal control and H_2_O_2_-treated sporozoites as the minimal signal control. The assay showed an excellent signal window (SW = 11.52) and Z′ factor (Z′ = 0.73), supporting its suitability for high-throughput screening (Fig. 2B; S2A Fig).

Primary screening of 5,000 bioactive compounds at 40 μM identified 28 compounds with ≥50% inhibition of sporozoite viability, including 14 compounds with >60% inhibition, 7 with >75% inhibition, and 3 with >90% inhibition (Fig. 2B; S2B Fig; S1 Table). The 14 compounds showing >60% inhibition were then evaluated in a secondary screen at 4 μM. Five compounds retained >50% inhibition at this lower concentration: ZL0420 (80.6%), sulbactam (77.0%), abexinostat (68.9%), kojic acid (57.0%), and SIB-1757 (62.1%) (Fig. 2C; S2 Table).

Dose-response assays confirmed that all five compounds inhibited sporozoite viability with submicromolar 50% effective concentrations (EC_50_ values) under the 3 h assay condition (Fig. 3A and 3B). Sulbactam was the most potent against free sporozoites (EC_50_ = 0.073 μM), followed by ZL0420 (0.091 μM), SIB-1757 (0.153 μM), abexinostat (0.189 μM), and kojic acid (0.311 μM). Consistent with their effects on ATP-based viability, treatment with these compounds at 10 μM induced visible morphological changes in sporozoites over time (Fig. 4). DMSO-treated sporozoites retained their typical curved morphology, whereas compound-treated sporozoites showed progressive deformation, becoming most apparent after 1–2 h of treatment.

**Fig 3.**
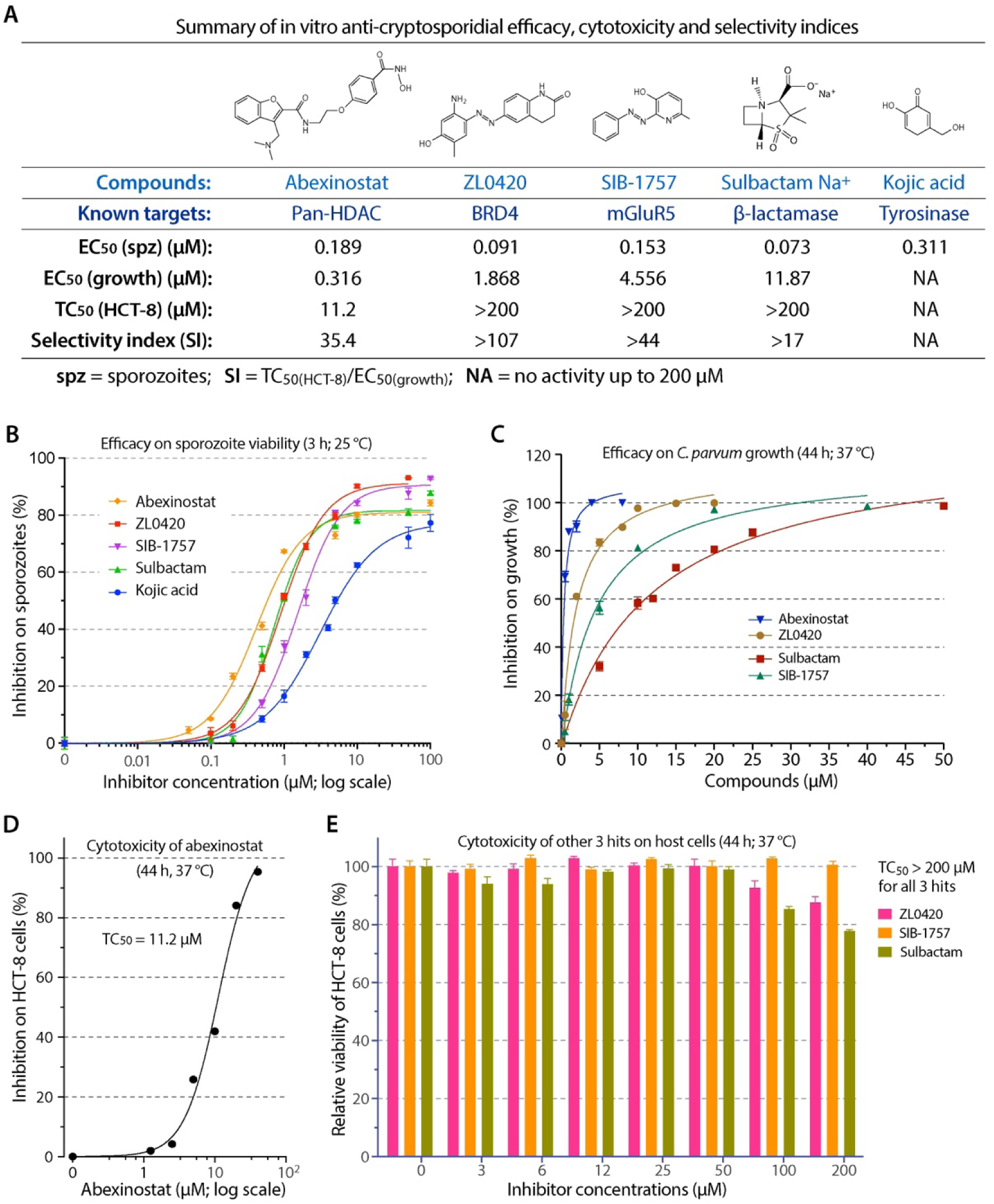
In vitro efficacies and cytotoxicity of the five top hits on *C. parvum*. **(A)** Summary of the structures of the five hits (abexinostat, ZL0420, SIB-1757, sulbactam, and kojic acid), their known molecular targets, 50% effective concentrations (EC_50_ values) on post-excystation sporozoites (spz) and parasite intracellular development (growth), and 50% toxic concentration (cytotoxicity; TC_50_ values) on HCT-8 cells by MTS assay. In vitro selectivity index (SI) values were calculated as the ratio between TC_50_ and EC_50_ (SI = TC_50(HCT8)_/EC_50(growth)_). **(B)** Dose-response efficacy curves of the five hits on sporozoite viability by luminescence ATP assay. Post-excystation sporozoites received 3 h treatment with serially diluted compounds at 25 °C, and evaluated by luminescence ATP assay. **(C)** Dose-response efficacy curves of four hits on the parasite intracellular development by the 44-h infection/qRT-PCR assay. In this assay, kojic acid showed no inhibition on the parasite growth at up to 100 μM. **(D–E)** Cytotoxicity of abexinostat (D) and the three other hits (ZL0420, SIB-1757, and sulbactam) (E) on HCT-8 cells (44 h assay).

**Fig 4.**
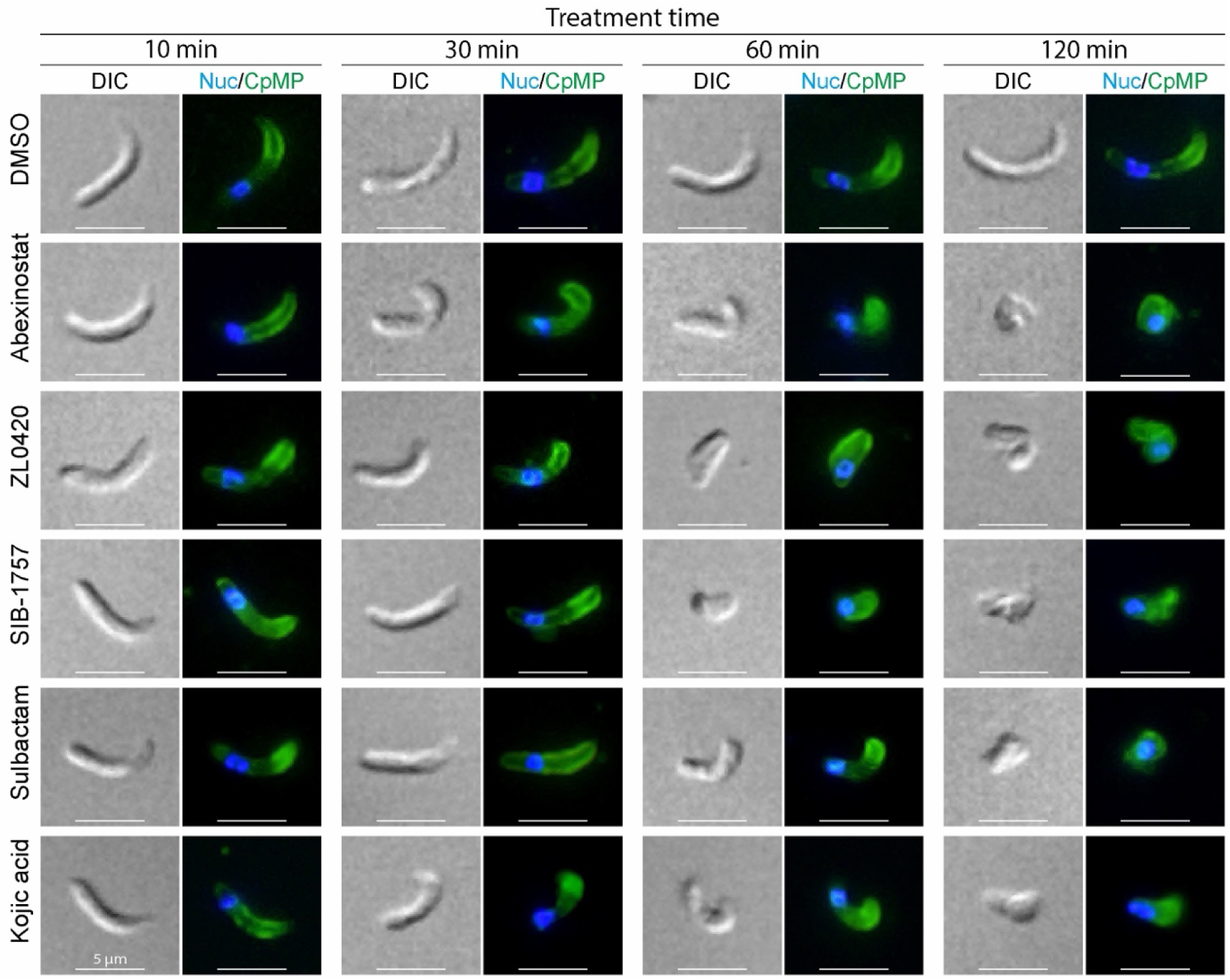
The morphology of *C. parvum* sporozoites in response to the treatment of five top hits. Post-excystation sporozoites were treated with abexinostat, ZL0420, SIB-1757, sulbactam, and kojic acid at 10 μM for up to 2 h. Samples were immunolabeled with a rabbit polyclonal antibody against a *C. parvum* membrane protein (CpMP) for immunofluorescence microscopy. Brightfield images were captured with differential interference microscopy (DIC). Compound-treated sporozoites show various degrees of deformation over time, while solvent (0.5% DMSO)-treated sporozoites retain normal morphology.

The five compounds have reported molecular targets or activities in other biological systems (Fig. 3A): abexinostat is a pan-HDAC inhibitor [47,48], ZL0420 is a BRD4 inhibitor [49], SIB-1757 is an mGluR5 antagonist [50,51], sulbactam is a β-lactamase inhibitor [52,53], and kojic acid is a tyrosinase inhibitor [54–56]. These annotations suggested that the sporozoite viability assay identified compounds associated with diverse biological activities. However, because the corresponding parasite targets may differ from their known mammalian or bacterial targets, direct biochemical validation was required for selected activities.

### Four hits inhibit intracellular *C. parvum* growth in vitro with favorable selectivity

We then evaluated the five sporozoite-active hits for inhibition of intracellular *C. parvum* growth using an established 44 h HCT-8 cell infection assay with parasite burden quantified by qRT-PCR. Four of the five compounds retained activity against intracellular parasite development (Fig. 3A and 3C). Abexinostat showed the highest potency in this assay, with an EC_50_ of 0.316 μM, followed by ZL0420 (1.868 μM), SIB-1757 (4.556 μM), and sulbactam (11.87 μM). Kojic acid showed no detectable inhibition of intracellular parasite growth at concentrations up to 200 μM.

Host-cell cytotoxicity was then evaluated in HCT-8 cells over the same 44 h exposure period. Abexinostat showed measurable cytotoxicity, with a 50% cytotoxicity concentration (TC_50_) of 11.2 μM, resulting in an in vitro selectivity index of 35.4 based on the ratio of TC_50_ in HCT-8 cells to EC_50_ against parasite growth (Fig. 3D). ZL0420, SIB-1757, and sulbactam showed minimal cytotoxicity at concentrations up to 200 μM, yielding selectivity indices of >107, >44, and >17, respectively (Fig. 3E).

Thus, four of the five compounds identified by the sporozoite viability screen also inhibited intracellular parasite development with favorable in vitro selectivity. For all four compounds, EC_50_ values were lower against free sporozoites than against intracellular parasite growth, consistent with the screen enriching for compounds that rapidly affect the invasive sporozoite stage.

### Four in vitro-active hits reduce *C. parvum* infection severity in IFN-γ-knockout mice

The four compounds with confirmed in vitro activity were further evaluated in IFN-γ-knockout mice infected with transgenic *C. parvum* expressing nanoluciferase with oocyst shedding quantified with nanoluciferase assay [19]. Infected mice were treated once daily for seven days beginning at 3 days post-infection (dpi). Based on available pharmacological information and previous in vivo use of these compounds [49,57–60], abexinostat was administered by oral gavage at 25 mg/kg/day, whereas ZL0420, sulbactam, and SIB-1757 were administered by intraperitoneal injection at 10, 20, and 20 mg/kg/day, respectively (Fig. 5A). These doses produced no overt adverse effects in uninfected C57BL/6 mice during a 7-day dosing period, based on body weight, fur condition, activity, posture, and mortality (S3 Table).

**Fig 5.**
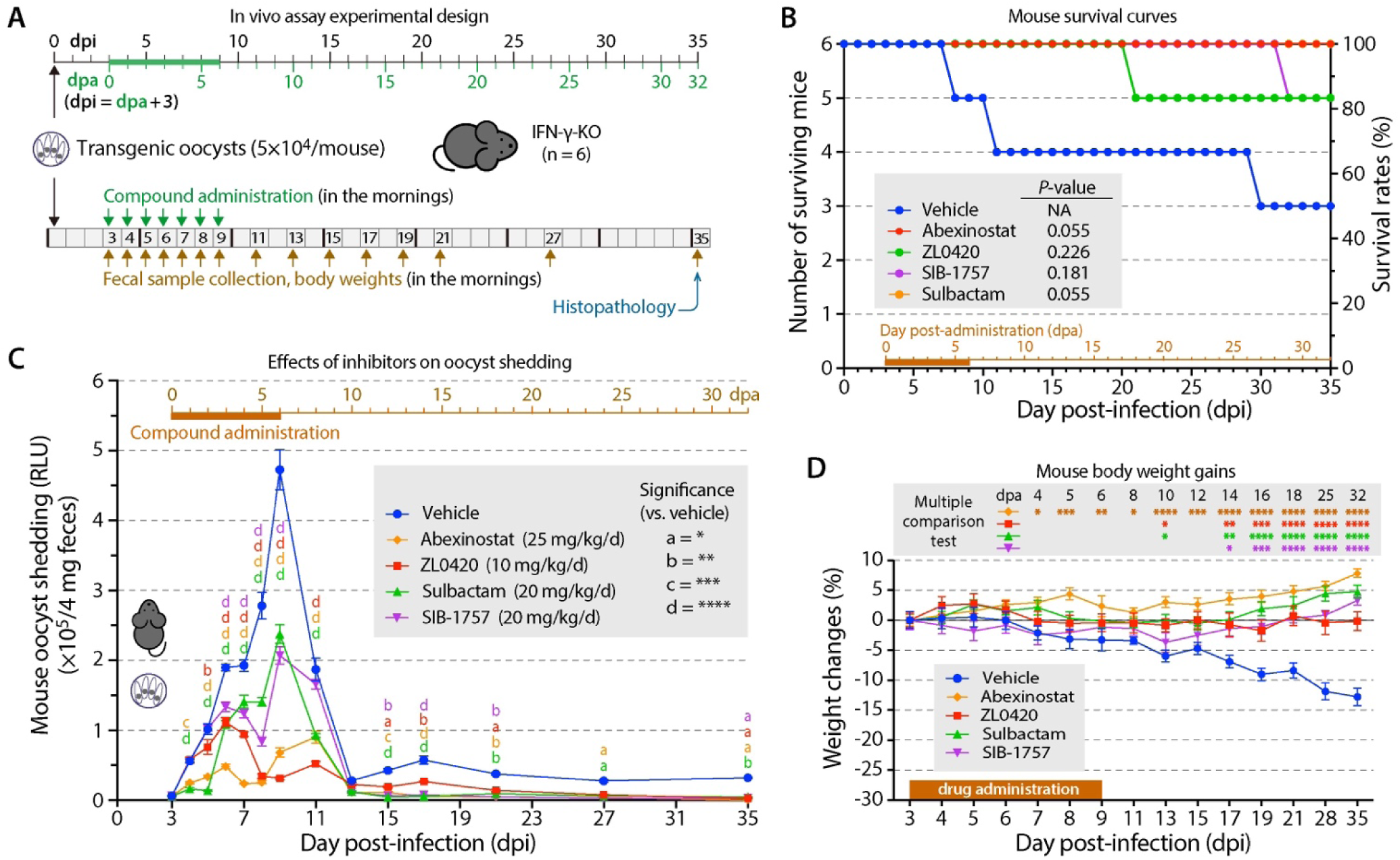
In vivo efficacies of abexinostat (25 mg/kg/d), ZL0420 (10 mg/kg/d), SIB-1757 (20 mg/kg/d), and sulbactam (20 mg/kg/d) against *C. parvum* infection in IFN-γ-knockout mice. (**A**) Schematic of the in vivo experimental design. Mice were inoculated with transgenic *C. parvum* oocysts expressing nanoluciferase (5 × 10^4^/mouse), allowed to establish infection for three days, and administered with compounds at specified single daily doses on 3 day post-infection (dpi) for seven consecutive days. Fecal samples were collected, and mouse body weights monitored on 3 dpi (= 0 dpa; day post-administration) and onward at specified intervals. Mice were sacrificed on 35 dpi (32 dpa) for histopathology. **(B)** Survival curves of mice treated with specified compounds during the course of infection. No deaths occurred in the abexinostat- or sulbactam-treated groups, while one death occurred in each of the ZL0420- and SIB-1757-treated groups. Survival differences did not reach statistical significance. (**C**) Oocyst shedding from mice, which peaked at 9 dpi in the vehicle group. All four compounds reduced oocyst shedding at multiple time points under the tested regimens. Statistical significances were calculated by Dunnett multiple comparison test versus vehicle, with significance levels marked in the graph. Also see Table 1 for area-under-the-curve data. **(D)** Mouse body weight gains/losses, showing steady weight decline in the vehicle group. All four compounds fully curbed the weight decline, with abexinostat and sulbactam groups showing significant weight gains.

**Table 1.** Oocyst shedding in mice as determined by nanoluciferase assay reported as relative luminescence unit (RLU)

| Treatment | Time period <sup>1</sup> | AUC <sup>2</sup> | SEM | Relative level | Reduction |
| --- | --- | --- | --- | --- | --- |
| Vehicle | Whole period | 2,734,778 | 141,577 | 100.0% | 0.0% |
|  | During treatment | 1,058,525 | 69,322 | 100.0% | 0.0% |
|  | After treatment | 1,676,253 | 123,444 | 100.0% | 0.0% |
| Abexinostat<br>(25 mg/kg/d) | Whole period | 574,389 | 52,976 | 21.0% | 79.0% |
|  | During treatment | 193,476 | 17,136 | 18.3% | 81.7% |
|  | After treatment | 380,913 | 50,128 | 22.7% | 77.3% |
| ZL0420<br>(10 mg/kg/d) | Whole period | 829,611 | 47,731 | 30.3% | 69.7% |
|  | During treatment | 393,786 | 33,057 | 37.2% | 62.8% |
|  | After treatment | 435,824 | 34,431 | 26.0% | 74.0% |
| Sulbactam<br>(20 mg/kg/d) | Whole period | 1,126,984 | 62,904 | 41.2% | 58.8% |
|  | During treatment | 542,415 | 38,139 | 51.2% | 48.8% |
|  | After treatment | 584,570 | 50,024 | 34.9% | 65.1% |
| SIB-1757<br>(20 mg/kg/d) | Whole period | 1,266,499 | 69,270 | 46.3% | 53.7% |
|  | During treatment | 608,230 | 38,531 | 57.5% | 42.5% |
|  | After treatment | 658,269 | 57,565 | 39.3% | 60.7% |

<sup>1</sup> Whole period = 3 to 35 dpi; During treatment = 3 to 9 dpi; After treatment = 9 to 35 dpi. <sup>2</sup> AUC, area under the curve.

In the infected vehicle group, three of six mice died during the 35-day experiment, with one death each on 8, 11, and 30 dpi (Fig. 5B). No deaths occurred in the abexinostat- or sulbactam-treated groups, while one death occurred in each of the ZL0420 and SIB-1757 groups. Thus, survival was numerically improved in all treatment groups, particularly in the abexinostat and sulbactam groups, although survival differences did not reach statistical significance under the group size used in this experiment.

Fecal oocyst shedding, measured by nanoluciferase activity, peaked at 9 dpi in the vehicle group (Fig. 5C). All four treatments reduced oocyst shedding during the treatment period and at multiple post-treatment time points. Area-under-the-curve analysis showed that, over the full 3– 35 dpi observation period, abexinostat reduced total oocyst shedding by 79.0%, followed by ZL0420 (69.7%), sulbactam (58.8%), and SIB-1757 (53.7%) (Table 1). During the treatment period, reductions were 81.7%, 62.8%, 48.8%, and 42.5%, respectively. During the post-treatment period, reductions were 77.3%, 74.0%, 65.1%, and 60.7%, respectively. Because the compounds were administered at different doses and by different routes, these values should be interpreted as efficacy under the tested regimens rather than direct rank-order comparisons of intrinsic in vivo potency.

Body-weight trajectories further supported treatment benefit (Fig. 5D). Vehicle-treated mice progressively lost weight, with average losses of 3.3% by 9 dpi and 12.8% by 35 dpi. All four treatments attenuated or prevented this weight loss. Abexinostat-treated mice gained weight throughout the experiment, with average increases of 2.4% at 9 dpi and 7.8% at 35 dpi.

Sulbactam- and SIB-1757-treated mice showed little change during the early phase but gained weight during the later phase of infection, while ZL0420-treated mice largely maintained body weight through the end of the experiment.

Together, these data indicate that the four in vitro-active hits reduced infection severity in vivo, as reflected by decreased oocyst shedding and improved body-weight trajectories. Survival outcomes were also numerically improved, but should be interpreted cautiously because statistical significance was not reached.

### Treatments partially improve infection-associated ileal pathology

Ileal tissues were collected at 35 dpi to assess the effect of treatment on intestinal pathology. In uninfected mice, the ileal mucosa showed elongated villi with preserved epithelial architecture (Fig. 6A). In infected vehicle-treated mice, the ileum showed marked villus shortening and epithelial disruption, with numerous epicellular parasites visible along the villus surface (Fig. 6B). In mice treated with abexinostat, ZL0420, SIB-1757, or sulbactam, ileal architecture was partially improved compared with vehicle-treated infected mice, although the tissue was not fully restored to the appearance of uninfected controls (Fig. 6C–F).

**Fig 6.**
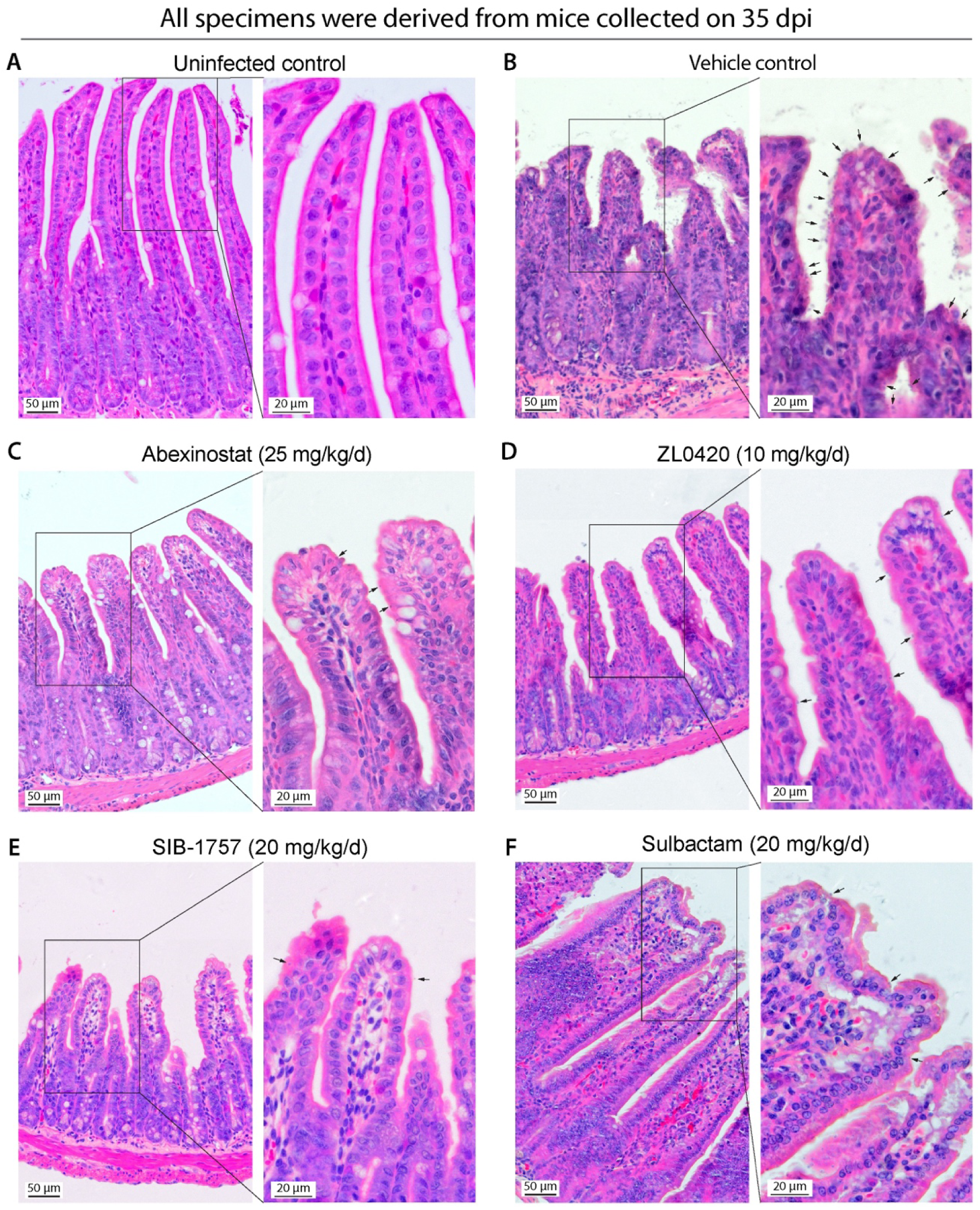
Histopathology of ileal tissues from mice treated with specified compounds and collected on 35 dpi. **(A)** Ileal tissues from an uninfected mouse showing elongated villi with smooth epithelial architecture. **(B)** Ileal tissues from an infected/vehicle control mouse, showing severely damaged epithelium with shortened villi with rough surface and numerous epicellular *C. parvum* (marked with arrows). **(C–F)** Ileal tissues from infected mice treated with abexinostat (C), ZL0420 (D), SIB-1757 (E), and sulbactam (F), showing partial improvement of villus architecture and epithelial morphology. Also see Fig 7 for the measurements of villus/crypt ratios.

Quantitative morphometric analysis supported the histological observations (Fig. 7). The ileal villus length/crypt depth ratio was markedly reduced in infected vehicle-treated mice compared with uninfected mice (P < 0.0001 between infected/vehicle and uninfected). Treatment with abexinostat, ZL0420, SIB-1757, or sulbactam increased the villus length/crypt depth ratio relative to the vehicle group, indicating partial recovery of ileal architecture. Because measurements were nested within individual mice, statistical comparisons were performed using nested analysis with mouse as the biological unit. Although all four treatments improved villus/crypt ratios relative to vehicle-treated infected mice at various significance levels (P values between <0.01 and <0.0001), the ratios remained below those of uninfected mice (P values between <0.01 and <0.0001), indicating partial but incomplete histological recovery by 35 dpi.

**Fig 7.**
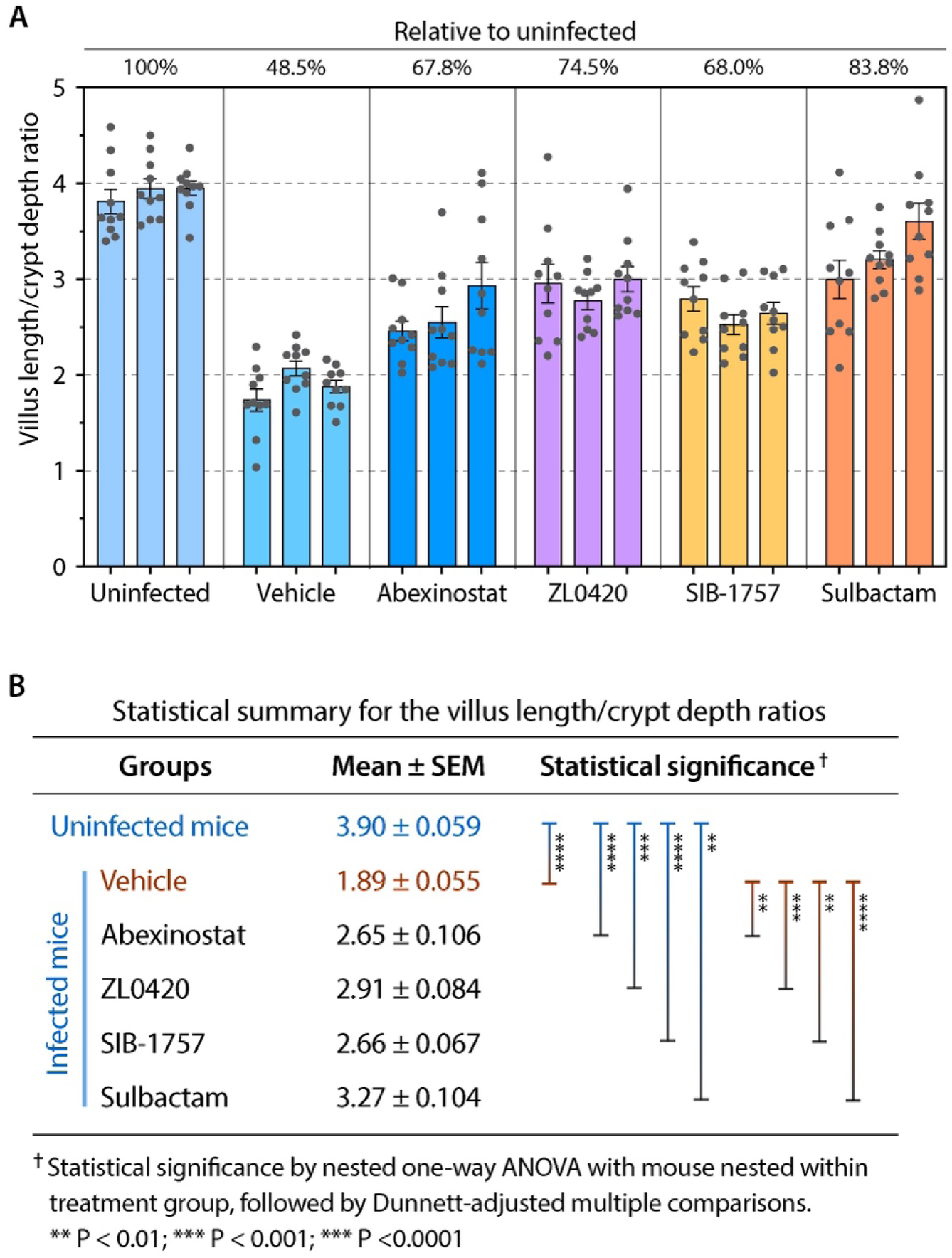
Ileal villus length/crypt depth ratios in mice treated with specified compounds and collected on 35 dpi. **(A)** Comparison of ileal villus length/crypt depth ratios between uninfected healthy mice and *C. parvum*– infected mice receiving vehicle, abexinostat, ZL0420, SIB-1757 and sulbactam. **(B)** Statistical summary of the ileal villus length/crypt depth ratios between uninfected and infected mice receiving various treatments. Each point represents one villus–crypt unit; 10 villus–crypt units were measured from each of three mice per group. Statistical comparisons were performed using nested one-way ANOVA, with mouse nested within treatment group, followed by Dunnett-adjusted multiple comparisons.

These histopathological and morphometric data are consistent with the oocyst-shedding and body-weight results, supporting in vivo efficacy of the four compounds under the tested regimens.

### Abexinostat and sulbactam inhibit native enzymatic activities in sporozoite lysates

Because abexinostat and sulbactam are annotated as inhibitors of HDAC and β-lactamase, respectively, we next tested whether corresponding enzymatic activities could be detected in *C. parvum* sporozoite lysates and inhibited by these compounds. We focused on these two activities because HDACs are encoded by *C. parvum* and were previously implicated as anti-cryptosporidial drug targets, whereas β-lactamase-like activity could be directly assayed using nitrocefin hydrolysis.

*C. parvum* encodes four annotated HDAC/Sir2-family proteins: two class I HDACs, one ankyrin-repeat-containing class II HDAC, and one class III/Sir2-family HDAC [61,62]. In contrast, the parasite lacks canonical orthologs of bacterial-type β-lactamases and mammalian BRD4, but encodes several metallo-β-lactamase or metallo-β-lactamase domain-containing proteins and multiple bromodomain-containing proteins. No obvious mGluR5 ortholog was identified. These genomic observations suggest that the parasite targets of sulbactam, ZL0420, and SIB-1757 may differ from the canonical targets known in bacteria or mammals.

Using recombinant *E. coli* β-lactamase as a positive control, sulbactam inhibited nitrocefin-hydrolyzing activity with an IC_50_ (50% inhibition concentration) of 5.66 μM and a Hill slope of 1.15 (Fig. 8A). Sulbactam also inhibited nitrocefin-hydrolyzing activity in crude *E. coli* lysates and *C. parvum* sporozoite lysates, with IC_50_ values of 19.64 μM and 41.24 μM, respectively (Fig. 8B). The corresponding Hill slopes were 1.24 and 1.29. These data support the presence of a sulbactam-inhibitable β-lactamase-like activity in sporozoite lysates, although the responsible parasite protein remains to be identified.

**Fig 8.**
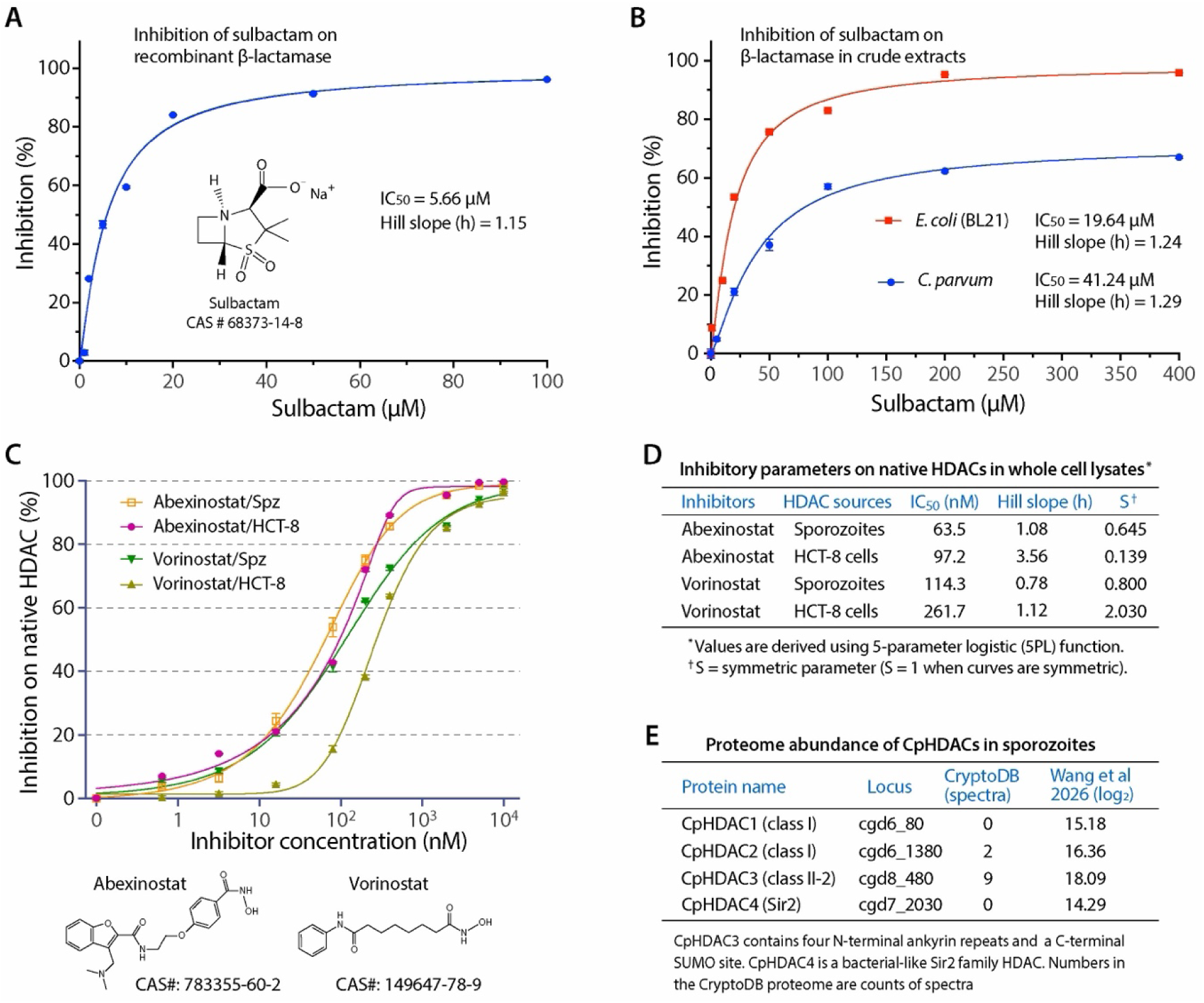
Inhibition of native enzymatic activities by sulbactam and abexinostat in *C. parvum* sporozoite lysates. **(A–B)** Sulbactam exhibited low micromolar activity on recombinant *Escherichia coli* β–lactamase as an assay control (IC_50_ = 5.66 μM) (A), and on the crude extracts of *E. coli* cells (IC_50_ = 19.64) and *C. parvum* sporozoites (IC_50_ = 41.24 μM) (B), confirming the presence of sulbactam-inhibitable β–lactamase–like activity in the parasite. The inhibitory kinetics were modeled by a four-parameter logistic curve fit, showing negligible cooperativity with Hill slopes ranging from 1.15 to 1.29. **(C–D)** Inhibitory curves (C) and summary of the kinetic parameters (D) for the two HDAC inhibitors (abexinostat and vorinostat) on the crude extracts of *C. parvum* sporozoites and HCT-8 cells. The kinetic curves were modeled with five-parameter logistic curve fit, showing low nanomolar IC_50_ values and varied Hill slopes and asymmetries. In this assay, vorinostat and HCT-8 cells were included for comparison. **(E)** Protein abundances of the four HDACs in *C. parvum* sporozoites. Spectral-count information was obtained from available CryptoDB proteomic data, and quantitative abundance values were obtained from our recently published *C. parvum* sporozoite proteome/phosphoproteome dataset. Protein abundance in the latter dataset was expressed as log_2_-transformed abundance values. The datasets confirmed the highest abundance of CpHDAC3, followed by CpHDAC2, CpHDAC1 and CpHDAC4. HDAC, histone deacetylase; IC_50_ value, 50% inhibitory concentration.

We also evaluated inhibition of native HDAC activity in sporozoite lysates by abexinostat, using vorinostat as a reference HDAC inhibitor. HCT-8 cell lysates were included for comparison.

Both abexinostat and vorinostat inhibited native HDAC activity in sporozoite and host-cell lysates with nanomolar potency (Fig. 8C and 8D). Abexinostat inhibited sporozoite HDAC activity with an IC_50_ of 63.5 nM and HCT-8 HDAC activity with an IC_50_ of 97.2 nM. Vorinostat showed IC_50_ values of 114.3 nM and 261.7 nM against sporozoite and HCT-8 HDAC activities, respectively. The IC_50_ value of vorinostat against sporozoite HDAC activity was comparable to our previous cell-based measurement of native parasite HDAC inhibition [61], and to its reported inhibition of recombinant CpHDAC3 [62].

Proteomic data further supported the presence of HDAC proteins in sporozoites. Among the four annotated CpHDACs, CpHDAC3 showed the highest abundance in both existing CryptoDB proteomic data and our recently published sporozoite proteome/phosphoproteome dataset, followed by CpHDAC2, CpHDAC1, and CpHDAC4 (Fig. 8E). These data do not identify which HDAC isoform is inhibited by abexinostat in sporozoite lysates, but they support native parasite HDAC activity as a targetable enzymatic activity in the invasive stage.

Together, the biochemical assays support inhibition of two native enzymatic activities in sporozoite lysates: HDAC activity by abexinostat and β-lactamase-like activity by sulbactam. These findings strengthen the biological interpretation of the sporozoite viability screen while indicating that the precise parasite protein targets, particularly for sulbactam, ZL0420, and SIB-1757, require further identification.

## Discussion

Phenotypic screening is one of the major approaches for discovering drug candidates against infectious diseases, including parasitic diseases [13,63–65]. For intracellular pathogens, however, in vitro screening is complicated by the need to culture pathogens in host cells and to quantitatively distinguish pathogen-derived signals from host-cell background. For C. parvum, several phenotypic assays have been developed to quantify intracellular parasite growth in epithelial cell monolayers, including qRT-PCR using cell lysates as templates, nanoluciferase assays using transgenic parasites, and high-content imaging assays based on fluorescent labeling of the parasitophorous vacuole membrane [19–22]. These assays have been highly useful for anti-cryptosporidial drug discovery, but they generally require one to two days of intracellular parasite development before parasite burden can be quantified.

In this study, we developed a complementary host cell-free approach that directly measures compound-induced loss of viability in excysted *C. parvum* sporozoites. The approach takes advantage of the observation that freshly excysted sporozoites can remain viable for several hours under defined extracellular conditions, providing a relatively short but sufficient window to identify fast-acting compounds. Among the three readouts evaluated, the luminescence ATP assay provided the best balance of sensitivity, linear dynamic range, assay time, parasite input, and cost. The assay is not intended to replace intracellular growth assays, because it does not capture compounds that act only during intracellular development, replication, or sexual differentiation. Instead, it provides a rapid front-end screening strategy that enriches for compounds capable of compromising the invasive stage before host-cell entry.

Using this luminescence ATP assay, we screened 5,000 bioactive compounds and identified five hits that retained >50% inhibition of sporozoite viability at 4 μM: abexinostat, ZL0420, SIB-1757, sulbactam, and kojic acid. All five showed submicromolar activity against free sporozoites. Four of them, except kojic acid, also inhibited intracellular parasite growth in vitro and displayed favorable selectivity relative to HCT-8 host cells. These results support the rationale that compounds affecting free sporozoite viability can include inhibitors with activity against later infection in host-cell culture. However, the sporozoite EC_50_ values should not be interpreted as direct predictors of intracellular growth potency. Differences between the two assay systems, including parasite stage, exposure environment, host-cell presence, medium composition, compound access, and assay duration, may all affect apparent potency. The lower EC_50_ values observed in sporozoites suggest that the invasive stage is particularly sensitive to these compounds under the conditions tested, but do not exclude additional effects on intracellular stages.

The four in vitro-active compounds also reduced infection severity in IFN-γ-knockout mice. All four treatments reduced fecal oocyst shedding, improved body-weight trajectories, and partially ameliorated ileal histopathology. Based on AUC analysis of oocyst shedding over the full 35-day experiment, the treatments reduced total shedding by 53.7–79.0%. These in vivo data provide important validation for the screening pipeline, because they show that compounds selected from the sporozoite viability assay can retain activity in a whole-animal infection model. At the same time, comparisons among the four compounds should be interpreted cautiously. The compounds were administered at different doses and by different routes, with abexinostat given orally and the other three compounds by intraperitoneal injection. Therefore, the relative reductions in oocyst shedding reflect efficacy under the tested regimens rather than direct rank-order comparisons of intrinsic in vivo potency. Survival was numerically improved in the treatment groups, especially with abexinostat and sulbactam, but survival differences did not reach statistical significance under the group size used in this experiment.

Among the validated leads, abexinostat provides the strongest mechanistic connection to a previously established anti-Cryptosporidium target class. The HDAC findings have both drug-development and biological implications. *C. parvum* encodes only a small set of HDAC/Sir2-family proteins, and the ankyrin-repeat-containing CpHDAC3 is highly divergent from mammalian HDACs [62]. This divergence suggests that selective inhibition of parasite HDACs may be achievable. However, abexinostat and vorinostat are broad-spectrum HDAC inhibitors and also inhibit host-cell HDAC activity. Therefore, while these compounds are useful chemical probes and leads, further optimization should aim to improve parasite selectivity, intestinal exposure, and safety. Isoform-specific biochemical assays, structural studies, and genetic or chemical-genetic validation will be important to define the relative contributions of individual CpHDACs to parasite viability, intracellular development, and gene regulation.

Sulbactam represents a second mechanistically interesting lead. In bacteria, sulbactam acts as a β-lactamase inhibitor and is commonly used in combination with β-lactam antibiotics [52,53]. *C. parvum* lacks canonical bacterial-type β-lactamase orthologs, but encodes several metallo-β-lactamase or metallo-β-lactamase domain-containing proteins. In the present study, sulbactam inhibited nitrocefin-hydrolyzing activity in sporozoite lysates, supporting the presence of a sulbactam-sensitive β-lactamase-like activity in the parasite. However, this result should be interpreted as activity-based evidence rather than definitive target identification. The responsible parasite protein remains unknown, and the activity detected in crude lysates may reflect one or more enzymes with β-lactamase-like hydrolytic activity. Identification of the sulbactam-sensitive protein or proteins will be needed to determine whether this activity is directly responsible for the anti-sporozoite and anti-cryptosporidial effects of sulbactam.

The other two in vitro and in vivo-active compounds, ZL0420 and SIB-1757, further illustrate the value of phenotypic screening for revealing unexpected parasite vulnerabilities. ZL0420 is annotated as a BRD4 inhibitor in mammalian systems [49]. Although *C. parvum* lacks a canonical BRD4 ortholog, it encodes multiple bromodomain-containing proteins that could potentially be involved in chromatin-associated processes. Whether ZL0420 acts on one of these parasite bromodomain proteins or on an unrelated target remains to be determined. SIB-1757 is known as an mGluR5 antagonist in mammalian systems [50,51], but no obvious mGluR5 ortholog is present in *C. parvum*. Its anti-cryptosporidial activity therefore likely reflects an alternative, currently unknown mode of action. These observations are intriguing, but target assignment for ZL0420 and SIB-1757 will require additional studies, such as chemoproteomic profiling, resistant-mutant selection, thermal proteome profiling, affinity pulldown, or recombinant protein assays.

Several limitations of the present study should be noted. First, the screen was performed only against extracellular sporozoites. Therefore, compounds that act specifically on intracellular replication, merogony, gametogenesis, or host-cell-dependent processes may be missed. The sporozoite viability assay should therefore be viewed as complementary to existing intracellular growth assays rather than as a universal replacement. Second, the ATP-based readout measures metabolic viability and may not distinguish direct parasite killing from rapid metabolic inhibition. As with other luminescence-based assays, primary hits also require orthogonal confirmation to exclude assay interference. In this study, the major hits were supported by dose-response assays, sporozoite morphology, qRT-PCR-based intracellular growth assays, cytotoxicity testing, and mouse infection experiments, which together reduce this concern for the validated leads. Third, the in vivo experiments used an immunodeficient mouse model and non-equivalent dosing regimens. Additional pharmacokinetic, pharmacodynamic, toxicity, formulation, and dose-optimization studies will be needed before these compounds can be prioritized for therapeutic development.

Despite these limitations, the study demonstrates that sporozoite viability is a useful and biologically meaningful endpoint for anti-*Cryptosporidium* drug discovery. The assay is rapid, operationally simple, host cell-free, and compatible with high-throughput screening. It may be particularly useful as the first step in a tiered pipeline, in which fast-acting sporozoite-active compounds are rapidly identified and then advanced to intracellular growth assays, host-cell cytotoxicity testing, and animal models. In principle, similar strategies may also be adapted to other apicomplexan parasites for which sufficient numbers of purified extracellular invasive stages can be obtained, such as *Eimeria* sporozoites, *Toxoplasma* tachyzoites, or *Plasmodium* merozoites.

In summary, we developed a luminescence ATP-based sporozoite viability assay for rapid phenotypic screening of anti-*C. parvum* compounds. Screening of 5,000 bioactive compounds identified four leads with activity against free sporozoites, intracellular parasite growth, and mouse infection. Biochemical assays further supported inhibition of native parasite HDAC activity by abexinostat and β-lactamase-like activity by sulbactam. These findings establish sporozoite viability as a practical screening endpoint, identify new anti-*Cryptosporidium* leads for further optimization, and highlight targetable enzymatic activities in the invasive stage of the parasite.

## Data availability statement

All data underlying the findings described in this manuscript are provided within the manuscript and its Supporting Information files. The primary and secondary screening datasets are provided as S1 and S2 Tables, mouse short-term tolerability data are provided as S3 Table, and the source numerical data used to generate the main and supporting figures are provided as S4 Data. Previously published sporozoite proteomic datasets used for CpHDAC abundance analysis are cited in the manuscript.

## Funding

This work was supported by grants from the National Key Research and Development Program of China (award number 2022YFD1800200) and the Science and Technology Development Plan Project of Jilin Province (SKL202502005JC). The funders had no role in study design, data collection and analysis, decision to publish, or preparation of the manuscript.

## Competing Interests

The authors have declared that no competing interests exist.

## Supporting Information

**S1 Table. Primary screening of 5,000 bioactive compounds at 40 μM.** Chemical information for the compounds and % inhibition on *C. parvum* sporozoite viability are shown.

**S2 Table. Secondary screening results of 14 top hits at 4 μM.**

**S3 Table. Short-term tolerability evaluation of the four hit compounds in C57BL/6 mice.** Body weights were recorded daily. Health scores were assessed based on physical activity, fur condition, body posture, and mental state using the indicated scoring criteria.

**S4 Data. Source numerical data used to generate Figs 1–8 and S1–S2 Figs.**

**S1 Fig. Optimization of the luminescence ATP assay for sporozoite viability.** Conditions affecting the luminescence readouts including the freeze-thaw cycles (A), in which intact oocysts and excysted sporozoites were compared; shaking time before readout (B), DMSO concentrations (C), and the incubation time after the addition of substrate (D). The reliability of the assay was also evaluated by plotting the linearity between luminescence signals and the percent of live sporozoites spiked with ATP-depleted sporozoites (E). RLU, relative luminescence signals.

**S2 Fig. Quality assessment of the sporozoite viability-based high-throughput screening (HTS) assay.** (A) Plot of luminescence signals (RLU) in plate uniformity assay. Maximal signals (Max) and minimal signals (Min) were derived from wells containing 0.5% DMSO and 1.0 mM H2O2, respectively. (B) Plot of the primary screening result. A total of 5,000 bioactive compounds at 40 μM were screened in duplicates.

## Author contributions

**Conceptualization:** Peng Jiang, Dongqiang Wang, Guan Zhu.

**Data curation:** Peng Jiang, Dongqiang Wang.

**Formal analysis:** Peng Jiang, Dongqiang Wang, Guan Zhu.

**Funding acquisition:** Jigang Yin, Guan Zhu.

**Investigation:** Peng Jiang, Dongqiang Wang, Yiming Wang.

**Methodology:** Peng Jiang, Dongqiang Wang, Guan Zhu.

**Supervision:** Jigang Yin, Guan Zhu.

**Validation:** Dongqiang Wang, Guan Zhu.

**Visualization:** Peng Jiang, Dongqiang Wang, Guan Zhu.

**Writing – original draft:** Peng Jiang, Guan Zhu.

**Writing – review & editing:** Peng Jiang, Dongqiang Wang, Guan Zhu.

## Notes

### Competing Interest Statement

The authors have declared no competing interest.

